# Historical remote sensing highlights long-term persistence of Emperor Penguin (*Aptenodytes forsteri*) colonies

**DOI:** 10.1101/2025.07.31.667967

**Authors:** Martynas Bielinis, Michelle LaRue, Benjamin M. Kraemer, Catalina Munteanu

**Affiliations:** Department of Forest and Wildlife Ecology, University of Freiburg, Stefan Meier Str 76, Freibung im Breisgau 79104, Germany; Department of Physical Geography and Ecosystem Science, Lund University, Sölvegatan 12, S-223 62 Lund, Sweden; Faculty of Silviculture and Forest Engineering, Transilvania University of Brasov, Str. Şirul Beethoven 1, 500123 Brasov, Romania; School of Earth and Environment, University of Canterbury, Christchurch, New Zealand 8041; Freiburg Institute of Advanced Studies, Freiburg im Breisgau 79104, Germany

**Keywords:** historical remote sensing, Keyhole, Landsat, Sentinel, guano stain, Cape Washington

## Abstract

Historical remote sensing imagery, including digital Landsat and analog Keyhole have potential to inform Antarctic conservation by providing insights into habitat and population dynamics. A case in point is that their long temporal range, extending as far back as the 1960s, offers an unparalleled look into the distribution of Emperor Penguin (*Aptenodytes forsteri*) colonies and their habitat. Here we demonstrate that Keyhole and Landsat sensors are capable of detecting presence and long-term change in penguin guano on sea-ice despite challenging environmental conditions. For 18 of the 66 known emperor penguin colonies, we confirmed their presence in images that predate earliest published records. We further used dense time series from 1960s to 2024 of the Cape Washington colony to exemplify change in guano area over time - but found little variation in guano area. We show that the guano area undergoes two distinct stages during a breeding season (a stable and clustered early stage, and a diffused and widespread late stage). We correlated guano area with penguin colony size and found that guano area correlates with colony size (Spearman’s ρ = 0.59, *p*-value = 0.017). Taken together, our results suggest long term resilience of penguin colonies in face of global change, and highlight that the use of historical remote sensing imagery across the Antarctic can inform conservation efforts and benefit the ongoing historical studies of penguin colony dynamics. We highlight the benefit of remote sensing for documenting otherwise inaccessible colonies and for quantifying and understanding historical colony dynamics.

## Introduction

Emperor penguins *Aptenodytes forsteri* are the largest living penguin species (Le Maho, 1977; Stonehouse, 1953), and the only animal worldwide adapted to breeding on sea ice during polar winters (Pinshow & Welch, 1980). Emperor penguin survival and breeding success is dependent on persistence of a stable platform - typically sea ice, ice shelves or land at the colony site (Wienecke, 2015). Further, they require close access to open water (Le Scornec et al., 2025; Șen et al., 2025) to forage efficiently (Massom et al., 2009). Emperor penguins are philopatric – returning to the same breeding sites every year (Pearce, 2007) because they prefer consistent and predictable conditions (Labrousse et al., 2023). For instance, variability in the phenology of sea ice formation may cause stress to birds (Fretwell et al. 2014) - increased and consistent stress may have a negative influence on survival (Jouventin, 1975). The sea ice must remain stable while the penguins lay and incubate their eggs (Wienecke, 2015), while the chicks hatch (Fretwell & Trathan, 2021), and during créching (Le Bohec et al., 2005). Adults leave the colony to forage (Fretwell & Trathan, 2021; G. Kooyman et al., 2000) in December, before their weeks-long annual moult. During this time they lack waterproofing and hence also require stable sea ice or land (Wienecke, 2015). Although the species is adapted to the harsh conditions of the Antarctic, they are also dependent on specific environmental conditions during their long breeding and vulnerable moulting periods. But global change may affect the availability of sea ice (Eayrs et al., 2021), which in turn can impact food availability or juvenile survival and recruitment (Le Scornec et al., 2025). The species can respond to such changes immediately (Forcada & Trathan, 2009), or with a time delay (Jenouvrier et al., 2017; Trathan et al., 2020). This is why long-term monitoring of penguin colonies using dense time series is necessary to understand population dynamics.

Field records of emperor penguin colonies date back to the 19^th^ century, with the first colony described in 1844 (Wienecke, 2015). By 2015, 54 colonies were known (LaRue et al., 2015). Currently 66 emperor penguin colonies have been confirmed by field visits (2022/2023 breeding season edition of the Mapping Applications for Penguin Populations and Projected Dynamics project) (Humphries et al., 2017). While important for assessing characteristics such as penguin movement patterns, diet or behavior (Le Maho et al., 2011), field visits alone are costly and insufficient for assessing long term colony dynamics because they are limited in their coverage and biased towards accessible areas (Ancel et al., 2017). Colony monitoring based on remote sensing provides a low-cost alternative, higher revisit times for monitoring changes, and reduces direct stress to the colony (Kamenev, 1968). Additionally, historical remote sensing data dating back to the 1960s (EROS Center, 2017a) holds great potential to complement and temporally-extend other survey data (Munteanu et al., 2024).

Remote sensing advanced our understanding of emperor penguin ecology because colonies can be detected and enumerated using various satellite platforms (Barber-Meyer et al., 2007; Fretwell & Trathan, 2009). The discoveries of new colony locations (Fretwell & Trathan, 2009; M. A. LaRue et al., 2015; Trathan et al., 2020) not only increases our ability to monitor specific locations, but further informs fundamental ecological knowledge, for instance on intraspecific competition. Emperor penguin colonies remain evenly spaced - even with the doubling of colony locations since 2010 (Santora et al., 2020). Which is a typical signal of “territorial” behavior (Maher & Lott, 2000) and lends the idea that emperor penguins compete over resources amongst themselves - for prey but also for nesting space. Notably, review of high-resolution imagery provides fine-scale detail otherwise elusive to us, such as foraging pathways (e.g., direction of travel and exact location of ocean entry), and evidence of early ice breakouts (Fretwell et al., 2023). Finally, remote sensing provides evidence for species adaptability via colony “blinking” (interannual changes in colony presence (M. A. LaRue et al., 2015)) and within season relocation to seemingly more stable locations (i.e. ice shelves (Fretwell et al., 2014)). In the face of a changing Southern Ocean, knowing the behavioral ecology of the species across its entire distribution would facilitate detailed conservation action and decision-making.

Satellite-based remote sensing of emperor penguins has informed broad-scale research on the species in the past decade (Labrousse et al., 2022; M. LaRue et al., 2024), with key advances in our understanding of their behavioral ecology, but these analyses remain temporally limited. Because most emperor penguin colonies with longer-term datasets are in proximity to research stations, the vast majority of locations remain understudied. For example, in the 1960s only 10 colony locations were known, this number tripled by 2010. While remote sensing enabled the documentation of 66 locations with (presumably breeding) emperor penguins, there is only one location with long-term mark-recapture data (Pointe Geologie (Barbraud & Weimerskirch, 2001)). Thus, a major gap in our understanding is the longevity and persistence of these colony locations. Although recently discovered, it remains unclear if they have always been there or if colony locations are ephemeral, as the “blinking” behavior suggests. Historical remote sensing can add data to our spatial understanding of the species’ presence and inform the understanding of temporal changes of their presence (blinking or not), location (sea ice vs. ice shelf behaviors) and the environmental conditions (colony proximity to sea ice edge) for more than 60 years.

The presence and relative size of a guano stain on ice is key to monitoring colonies using remote sensing (Fretwell et al., 2015; M. A. LaRue, 2014). The guano composes a characteristic dark brown or orange stain on the white and featureless Antarctic fast ice (sea ice fastened to land), which makes it detectable in remote sensing imagery. Guano was used to locate penguin colonies remotely as far back as 1984, when spectral profiles of Adélie penguin guano were compared with those sensed remotely by Landsat Thematic Mapper (TM), Landsat Multispectral Scanner (MSS) and SPOT HRV sensors (Schwaller et al., 1984). While availability and analyses of modern medium and high resolution datasets is continuously expanding, historical remote sensing data remains underutilized for species monitoring (Munteanu et al., 2024). Two effective and relatively low-cost solutions for filling this temporal gap are represented by the historical Keyhole reconnaissance satellite program and the legacy Landsat data.

Historical remote sensing collected during the Cold War era for military reconnaissance purposes is now available for research in ecology and conservation (Cloud, 2001; Munteanu et al., 2024). Data from the Keyhole programme includes worldwide imagery from 1960 to 1984 (EROS Center, 2017a, 2017b, 2017c). The value of these datasets for environmental science has long been acknowledged, but research is mainly focused towards glacier ice analysis (Bolch et al., 2010; Narama et al., 2010; Zhou et al., 2018), land cover analysis (Barthelme et al., 2024; Chen et al., 2014; Kivinen & Kumpula, 2014; Mihai et al., 2016; Rizayeva et al., 2023), or archeology (Fowler & Fowler, 2005). Keyhole imagery has been used to monitor sea ice dynamics and its potential impact on penguins (Massom et al., 2009), but to our knowledge, no study to date employed these datasets to detect guano and infer colony dynamics.

Historical multispectral imagery provides several advantages for guano detection over panchromatic data (Keyhole). Guano reflects strongly in visible red and near-infrared (NIR) (Mustafa et al., 2017), and appears brown, red and pink in the visible light (Fretwell & Trathan, 2009). Guano is highly-reflective and distinctive in short-wave infrared (SWIR) (Fretwell et al., 2014; M. A. LaRue et al., 2015; Rees et al., 2017; Schwaller et al., 1984), but not spectrally-unique in visible light or NIR (Rees et al., 2017). This suggests that SWIR-carrying sensors are best used in guano detection. Furthermore, Sentinel-2 NIR (Fretwell & Trathan, 2021) or Landsat MSS NIR (Schwaller et al., 1984) are also capable of distinguishing guano. These data, and the fact that Landsat 4 MSS was used to identify Adélie penguin colony sites (Borowicz et al., 2018), indicate an unexplored potential for detecting historical colony presence and size using legacy Landsat images.

Beyond colony detection, remote sensing can provide estimates of colony size from very high resolution (VHR) imagery (∼30-50 cm spatial resolution), in which individual penguins can be identified (Labrousse et al., 2022), but guano signatures can also be used to derive approximate colony size - previous studies utilized Landsat, Sentinel and other coarser-resolution datasets (Fretwell et al., 2012; Fretwell & Trathan, 2021), and the area of Adélie penguin guano was shown to correlate with the size of the colony (M. A. LaRue, 2014). The uncertainty related to such estimates remains high given population fluctuations within and between breeding seasons (Fretwell et al., 2012; Kooyman & Ponganis, 2017). Moreover, at any given time, a portion of the colony is out hunting or moulting, and estimation itself can be challenging due to the sheer number and uneven density of the birds at the site (Wienecke, 2011). This is why combined datasets over longer time periods hold great potential to produce more reliable estimates of both colony presence and particularly long term temporal dynamics (Richter et al., 2018).

Our goal is to integrate historical and recent remote sensing data for tracking Emperor Penguin (*Aptenodytes forsteri*) presence and colony dynamics. To address this goal, we combined historical remote sensing data from different sensors, including optical imagery of the Keyhole missions from the Cold War era and Landsat legacy data for all known emperor penguin colonies in Antarctica, as well as modern Landsat and Sentinel data for the colony at Cape Washington. Our specific research questions are:

1. Can emperor penguin guano reliably and regularly be detected in the oldest historical remote sensing datasets going as far back as the 1960s? We expect that guano detection is possible using oldest available datasets (Keyhole satellite imagery of the Cold War period dating to the 1960s), but that detection will be dependent on environmental and weather conditions (e.g. time of year, sea ice extent, weather patterns), and the quality of remote sensing data (e.g., resolution, acquisition frequency) (M. LaRue et al., 2024; M. A. LaRue et al., 2011, 2017; Witharana & Lynch, 2016).
2. Can guano stains (i.e., colony presence) be confirmed across Antarctica for historical periods that predate first colony records? We expect that guano stains can be used as a reliable indication of colony presence and that the existence of contemporary colonies at the same location can be confirmed prior to existing remotely-sensed surveys.
3. Does guano area estimated from historical and contemporary remote sensing datasets correlate with known estimates of colony size? We expect that the size of the guano area correlates with colony size, and that guano dynamics can be tracked using remote sensing estimates of guano indices.

## Methods

### Study area

We studied 66 emperor penguin colony sites (S1, S2) across the coastline of the Antarctic continent, cross-checked against colony lists in (Fretwell & Trathan, 2021), and MAPPPD (Humphries et al., 2017). The coastline of Antarctica stretches between 63° south latitude at the tip of the Antarctic peninsula to 78-80° south latitude at various ice-free points near the Ross and Ronne ice shelves. At these extreme latitudes, conditions for remote sensing vary between the Antarctic winter (June – August) and the Antarctic summer (December – February) as daylight restricts optical sensing. Additionally, cloud cover is notably more frequent during the Antarctic summer months than during other seasons (Frey et al., 2018) (Figure 1). Imagery from September to mid-January was prioritized over imagery from February or March as lack of sea ice during the late austral summer and possibility of penguins leaving the colony at that time limit the colony detection (Wienecke, 2015).

**Figure 1.**
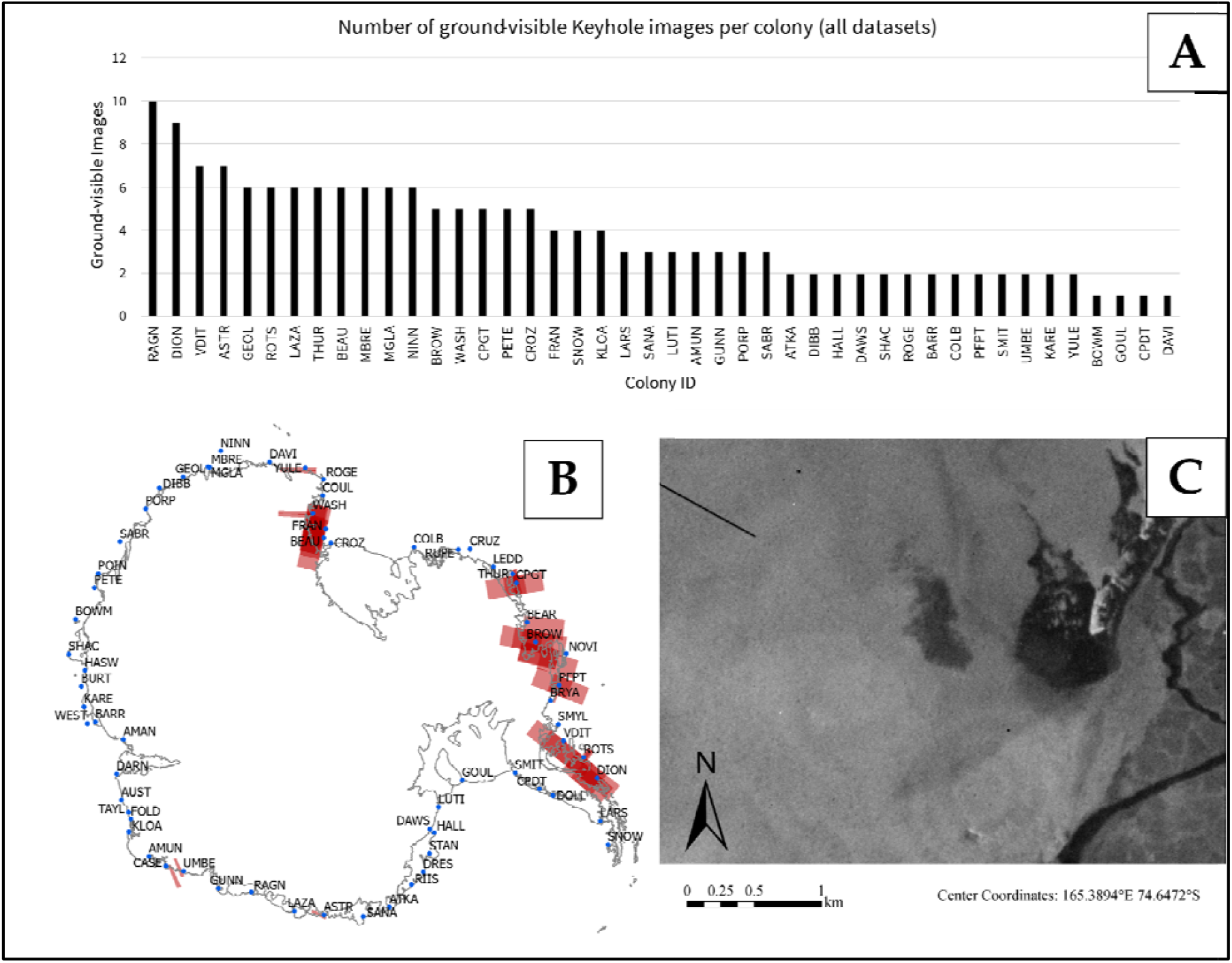
A: Number of utilized Keyhole images with ground visibility per colony site (22 colonies had no ground-visible Keyhole imagery); B: Declass-2 and Declass-3 Keyhole download-ready imagery spatial coverage. Antarctic land and ice shelf boundaries pictured in cyan (Gerrish et al., 2020); C: Visible guano at Cape Washington colony captured by KH-9 Gambit on September 22^nd^, 1980.

### Data

We used historical and modern remotely sensed imagery spanning the months of September to March for all years between 1960 and 2024 from Keyhole (S3), Landsat 1-7 (S4), Landsat 7-8, and Sentinel-2 (S4-5). Landsat 1-7 and Keyhole imagery (S6) were used for guano colony presence confirmation across the colony sites. Landsat 7-8 and Sentinel-2 imagery (S7) were used in addition for guano area measurement and subsequent correlation with colony size at Cape Washington.

Emperor penguin census data for the 66 colonies was obtained from the MAPPPD database^1^. The individual colony counts were published by (Kooyman & Ponganis, 2017; Korean Polar Research Institute, 2020; G. J. Wilson, 1983; Woehler, 1993).

We acquired scanned single-band panchromatic Keyhole (KH) imagery (S3) through the USGS EarthExplorer database^2^ (EROS Center, 2017a, 2017b, 2017c) as well as historical Landsat imagery (S4) taken between 1972-2003 (EROS Center, 1999, 2020a, 2020b). Thematic Mapper (TM) and Enhanced Thematic Mapper (ETM+) Landsat sensor imagery is offered with applied surface reflectance correction (U.S. Geological Survey, n.d.-a), while the raw Multispectral Scanner (MSS) data was corrected to top-of-atmosphere reflectance (U.S. Geological Survey, n.d.-b) given the negligible effect of the dry Antarctic atmosphere on the visual clarity of the imagery (Bindschadler et al., 2008). For colony size estimations, we relied on imagery from the Landsat 8 Operational Land Imager (OLI) and Sentinel-2 to cover the remaining time interval after the end of ETM+ operation (S5).

### Guano detection in historical optical imagery

For each colony location (S1), we applied a 25-kilometer search radius for imagery based on maximum observed coordinate offsets. The median location offset was ∼34.7 meters when compared to the study by (Fretwell & Trathan, 2021) (S8) and ∼36.7 when compared to MAPPPD (S9). Five colonies had large offsets (>1 km) compared to (Fretwell & Trathan, 2021), and 9 colonies had large offsets (>1 km) when compared to MAPPPD. Keyhole images were individually checked against their georeferenced footprints (as provided by USGS EarthExplorer) to ensure colony point presence within the scene.

For colony detection, we relied on download-ready Keyhole images. Landsat imagery was retrieved for each colony up to the ETM+ Scan-Line Corrector (SLC) fault date in 2003 (U.S. Geological Survey, 2003), after which ETM+ data contains characteristic data gaps. We acquired at least one clear acquisition per breeding season for TM/ETM+ sensor data, and all available imagery from MSS.

### Guano detection predating existing colony records

For all colonies, we considered the published colony date of discovery compiled by (Fretwell & Trathan, 2021). To verify if earlier colony presence predating the initial colony discovery date using remote-sensing data was available, we carried out an additional search for published studies with reported colony presence, using Clarivate™ (Web of Science™) and Google Scholar™ databases, using combinations of the following search string structures: “Colony Name”, “Colony Name” penguin*, “Colony Name” emperor penguin*, “Colony Name” remote sensing.

We recorded presence with those extracted from our time series of historical Keyhole, MSS, TM and ETM+. Cases in which presence is ambiguous or impossible to assess due to environmental and sensor conditions were excluded (S10). The earliest image dates begin in September/October, while the last breeding season imagery is from January or early February, when sea ice begins to melt at most colony sites. We did not record absences, because true absence cannot be assessed from the coarse resolution imagery, especially for smaller colonies.

#### Generalized Additive Modeling of Guano Area Trends

We used the Cape Washington colony (74.637° S 165.382° E) as a case study for assessing intrannual and interannual variation in guano patch size. Cape Washington has an overall stable size and long history of available imagery. In addition to legacy data, we used ETM+ (SLC-off data (U.S. Geological Survey, 2003)), Landsat 8 OLI and Sentinel-2 data with a 25-kilometer spatial filter. ETM+, OLI and Sentinel-2 data intake was limited to one image every ∼6 days (S7). We measured the guano area in visible-light conditions and short-wave infrared/near-infrared (SWIR/NIR), separately (S11). Because MSS lacks a visible blue band, the NIR band (Band 6) was included as a substitute for the visible-light. For Keyhole imagery we used the single panchromatic band. We manually traced and measured (square kilometers) guano area in each image.To characterize seasonal and long-term trends in penguin guano area, we fit a generalized additive model (Wood, 2017, 2023) with the mgcv^3^ package (version 1.9-1) in R (version 4.4.1). To account for potential biases from partial image obstruction (e.g., shadows and scan line corrector gaps), we calculated a potential error term as the sum of SWIR/VIS-specific shadow area and SLC error area.

Guano area measurements are subject to several sources of uncertainty that may limit their ability to precisely reflect colony size. One key factor is the temporal evolution of guano distribution over the breeding season. As the season progresses, penguin movement increases, leading to spatial dispersion of guano beyond the main nesting area. This results in a larger and more diffuse guano signature that may overestimate the actual colony size. Satellite images acquired later in the season (e.g., November) frequently show elongated guano tracks and scattered sub-clusters forming away from the central colony site (S12). In contrast, early-season images (September to October) typically reveal a denser and more spatially concentrated guano patch (S13), suggesting that guano area may more reliably represent colony size during this earlier, more stationary phase of the breeding cycle. Second, Landsat 7 ETM+ has known data gaps in imagery after May 31^st^, 2003 (U.S. Geological Survey, 2003) (S14). Existing corrections (Lee et al., 2016; Pringle et al., 2009) cannot retrieve missing data, therefore the dataset was analysed without these corrections. These data gaps were excluded from guano area measurements, and the maximum error area to which the guano patch could extend was calculated separately.

Observations were then weighted by the inverse of the percent error, downweighting highly uncertain values. The response variable (guano area in m^2^) was Yeo-Johnson transformed to improve normality. The GAM included a categorical predictor for data source (VIS or SWIR), a cyclic smoother for day of year, and a thin plate regression spline on decimal date, with smoothing parameters estimated using REML. All predictors were additive. We tested for interaction terms (e.g., by-source specific smooths for day of the year and decimal date) but found no evidence of source-specific trends (ΔAIC = +5.8), and thus retained a simpler additive model structure for interpretability.

### Correlation between guano stain size and colony size

To evaluate whether remotely sensed guano-covered area could serve as a proxy for penguin colony size, we first used the fitted GAM to generate daily predictions of guano extent for each year in the satellite record. Because the available colony size data - based on annual ground counts of chicks compiled in the MAPPPD database (Humphries et al., 2017) - were reported at annual resolution, we aggregated daily guano predictions into annual means to enable direct comparison. Guano area estimates were then merged with the MAPPPD colony size data to create a unified dataset, and we used Spearman’s rank correlation to quantify the association between guano area and colony size, reporting both the correlation coefficient and its associated p-value.

## Results

### Results of guano detection in historical optical imagery

For the 66 study colonies, we evaluated a total of 620 images for guano presences (144 Keyhole, 146 Landsat MSS, 128 Landsat TM and 202 Landsat ETM+). We identified guano in the oldest Landsat MSS imagery, where 6 out of 66 colonies have visible guano in the dataset, and this capability is further increased by TM and ETM+ sensors that have more frequent imaging and additional SWIR bands (guano visible in 24 and 33 colonies out of 66, respectively). Guano is also visible in Keyhole imagery, but visibility is dependent on colony size and the resolution of the dataset. With the exception of Cape Washington, we could not reliably identify guano in the freely available Keyhole imagery. Four colony locations were clearly visible in high-resolution Keyhole imagery (Thurston Glacier, Cape Gates, Beaufort and Brownson Island), but no guano presence was detected due to acquisition dates in March, during the Antarctic sea ice minimum (Turner et al., 2022). Although guano was often detected in Landsat, shortcomings such as ETM+ SLC-off data or high SWIR noise increased uncertainty estimates (S14-S16). Images from Keyhole are captured infrequently and irregularly (S17) and are not evenly distributed across the Antarctic – 36 colonies in high resolution Declass-3 and 39 colonies in medium resolution Declass-2 data have no imagery available (Figure 1, S18, S19),. This resulted in sparse temporal sampling that is affected by cloud cover, insufficient surface illumination or radiometric errors (S20).

### Results of guano detection predating existing colony records

Our analyses showed that historical remote sensing can be used to detect guano and infer presence at emperor penguin colonies. Importantly, for 18 of the analyzed colonies, our results are the earliest reported presences of the colonies (Figure 2, Figure 4, Table 1). To the best of our knowledge these represent the earliest observed guano patches predating colony discovery or other historical remote sensing studies on colony presence (Table 1). However, the total number of confirmed guano images is low, especially for the older MSS sensor (6 out of 22, Figure 2).

**Table 1:**
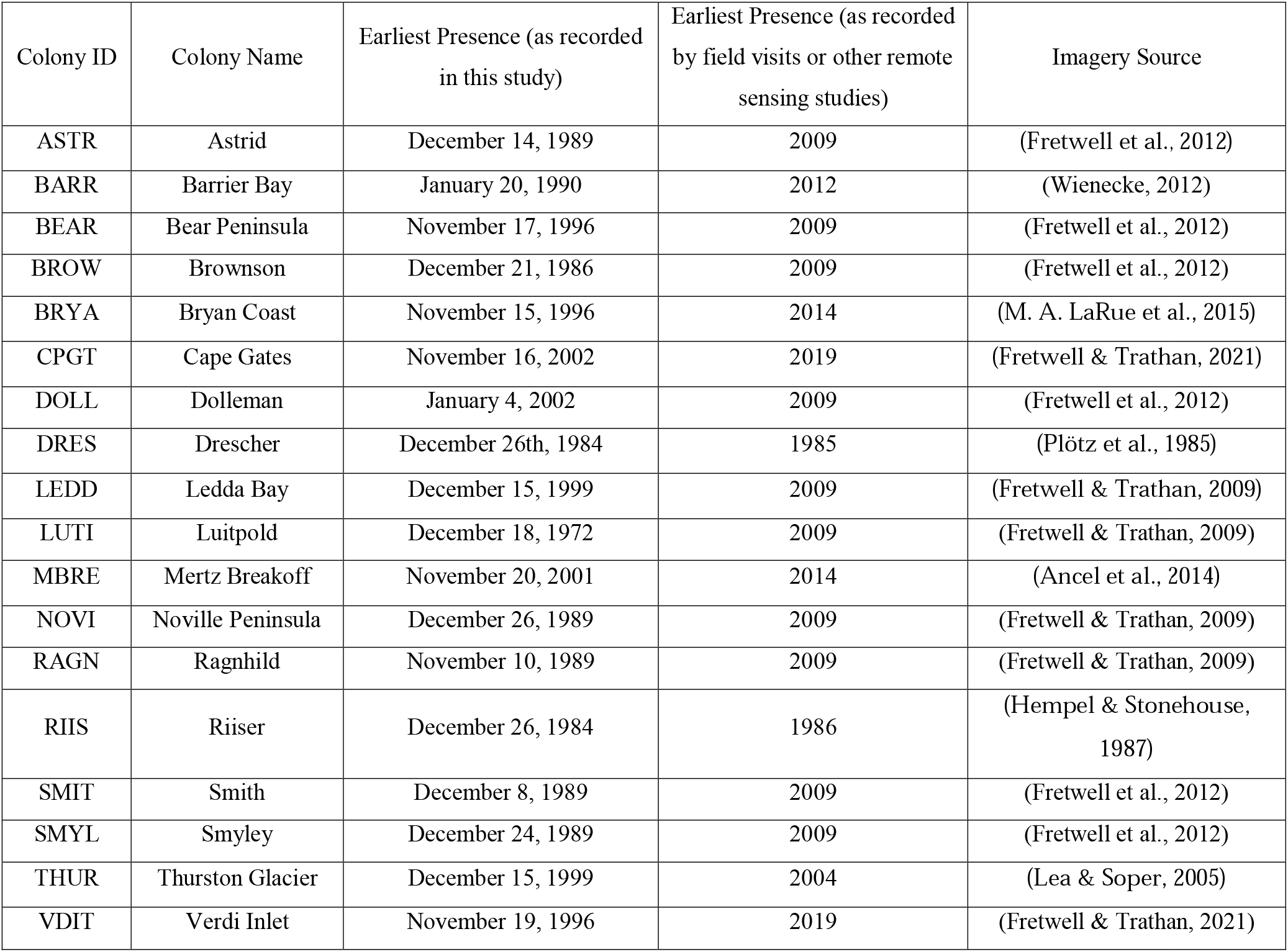
Colonies with analysed imagery predating oldest analyses imagery in studies (including date of earliest known imagery)

**Figure 2.**
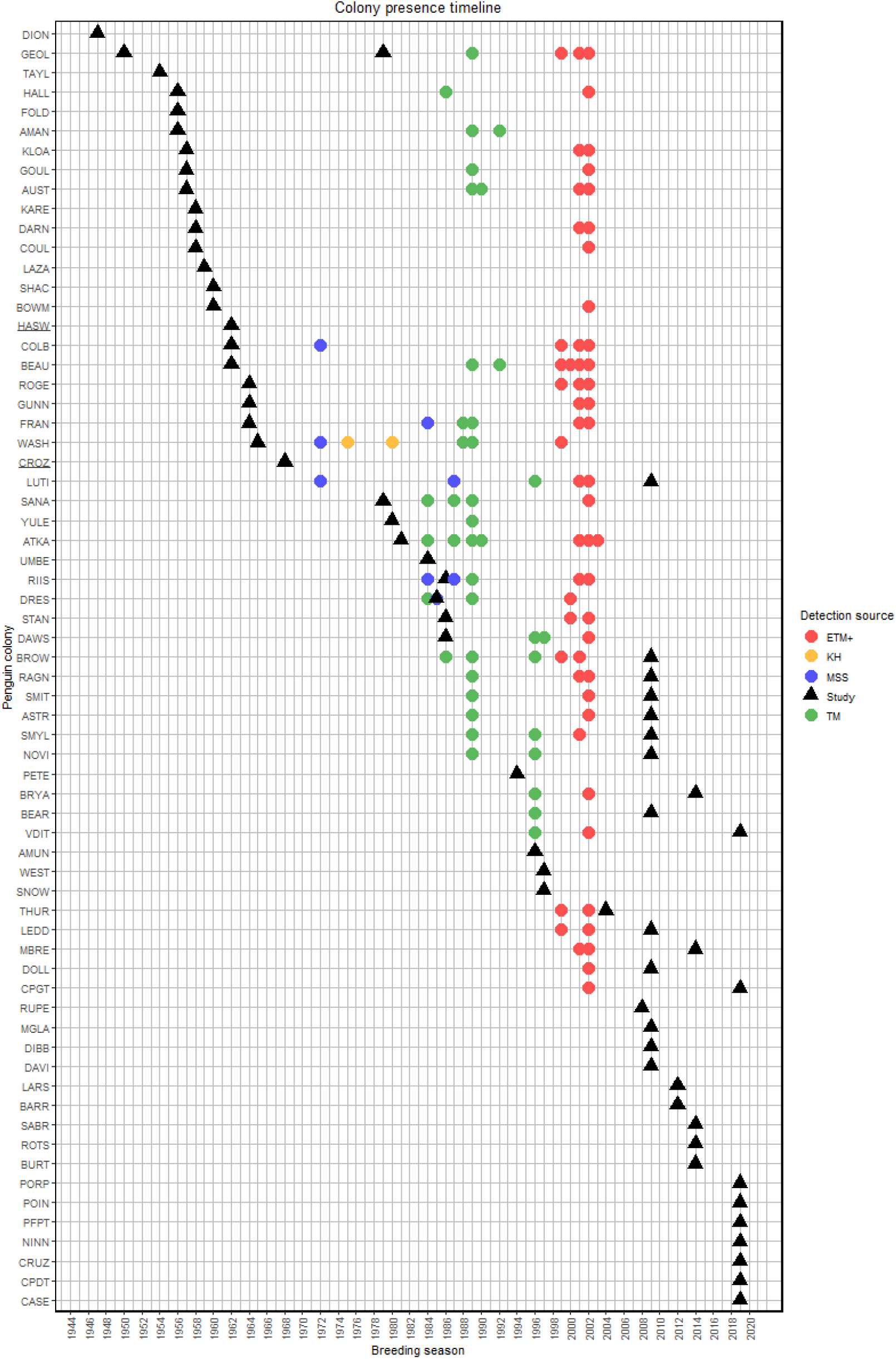
Detected guano (colony) presence by colony and breeding season. Note that for Cape Crozier (CROZ) and Haswell (HASW) colonies, the earliest reported presence exceeds the boundaries of the figure, and is 1902 (E. A. Wilson, 1907) and 1912 (Mawson, 1915), respectively.

### Results of correlation between guano stain size and colony size

For Cape Washington we analyzed 220 images between 1972 and 2024 for which we identified guano in 29 years. Overall, we found little variation between years, but rather substantial changes in guano patch size within a season. Despite an increase in visible-band guano between 2017 and 2024, the area stayed relatively stable over the observation period of 1972 and 2024 (Figure 3). The size of the observed guano patch in the visible band combination over the period of 29 years was on average 1 310 466 sq. m. (maximum area: 7 042 490 sq. m., minimum non-zero area: 42 538 sq. m.), while in the SWIR-NIR band combination it was on average 848 596 sq. m. (maximum area: 5 771 452 sq. m.; minimum non-zero area: 21 231 sq. m.).

**Figure 3.**
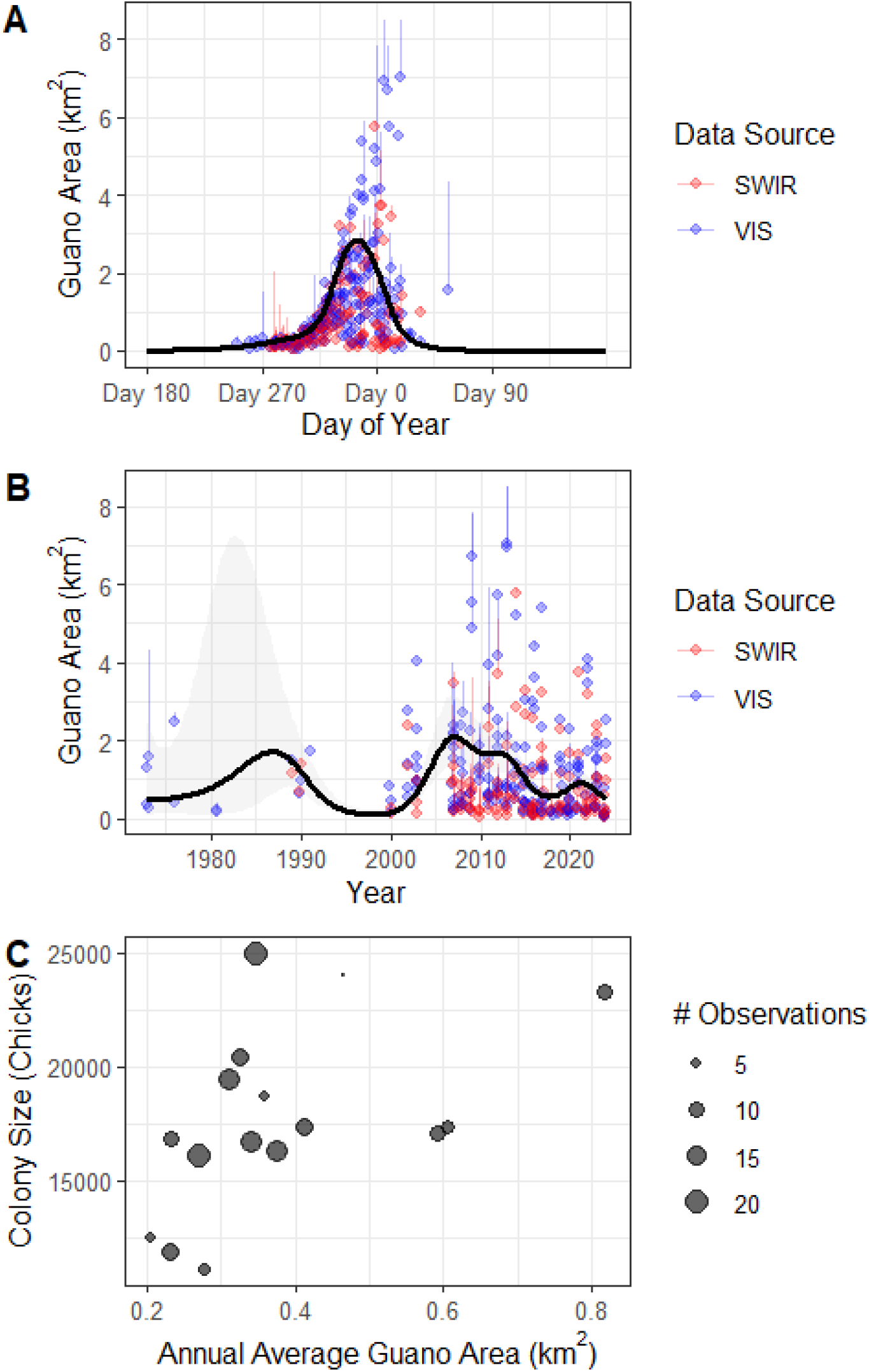
Seasonal and long-term variation in satellite-observed and modeled guano area, and its relationship with penguin colony size. (A) Intra-annual patterns of observed guano area (km^2^) across all available Landsat imagery using visible (VIS, blue) and shortwave infrared (SWIR, red) bands. Points represent the minimum confidently identified guano area, while erro bars extend to the maximum potential area including pixels likely obscured by shadows or Landsat SLC (scan line corrector) gaps. The black line shows the smoothed seasonal trend from a generalized additive model (GAM), predicted for the median year (2014). Day of year is centered on austral summer. (B) Long-term trend in guano area estimated from visible and SWIR imagery. Points show all individual observations with error bars indicating upper bounds due to shadow or SLC uncertainty. The black line shows the GAM-predicted multi-year trend with 95% confidence interval (gray ribbon), incorporating a shared seasonal component. (C) Relationship between annual average guano area (VIS-based, km^2^) and chick counts from ground-based colony surveys. Dot size reflects the number of satellite observations for that year. Years with few or uncertain observations are down-weighted, and some estimates rely only on modeled seasonal predictions. A significant positive correlation was found (Spearman’s ρ = 0.59, p = 0.017), suggesting that larger colonies tend to produce larger visible guano coverage.

**Figure 4.**
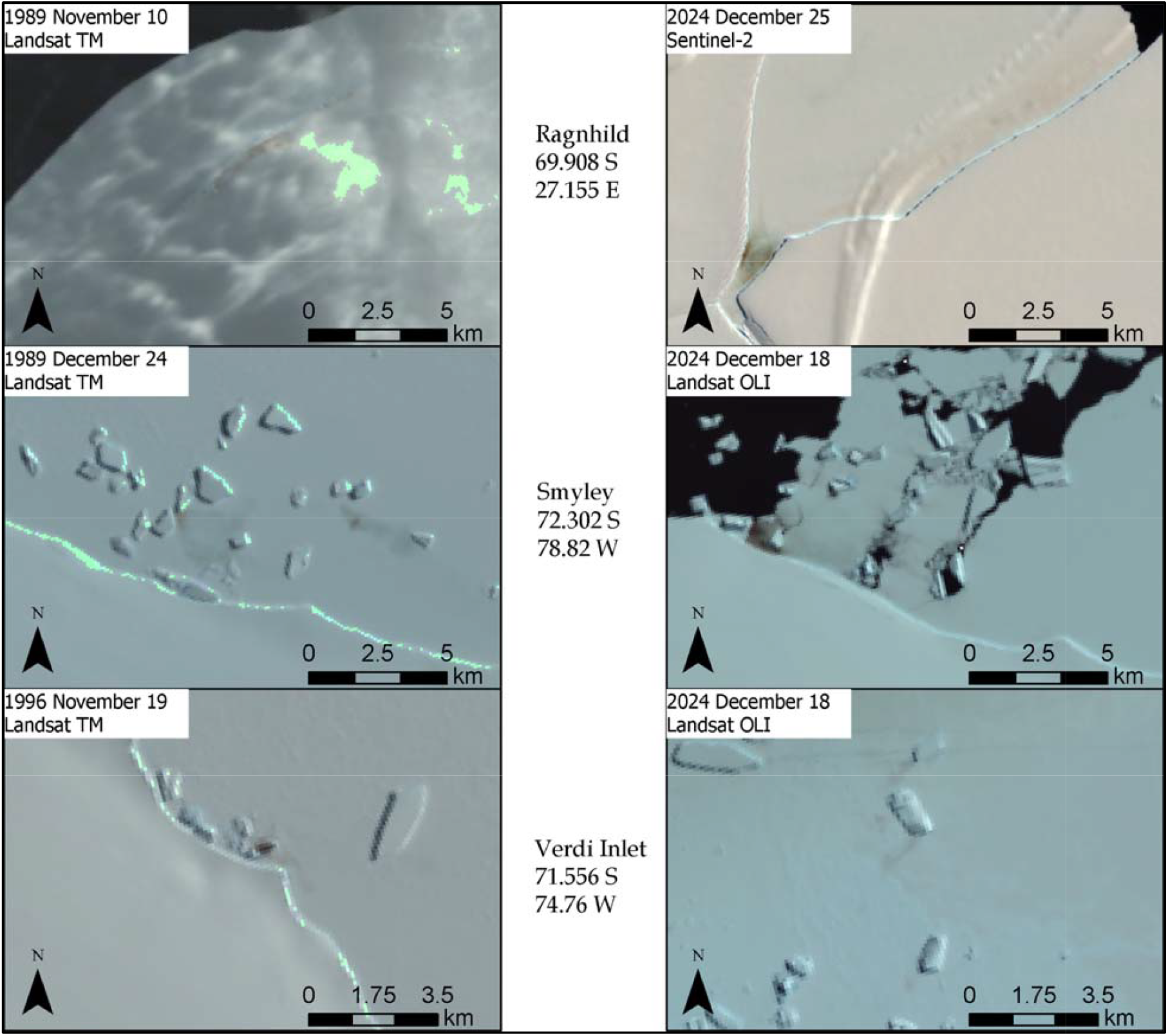
Imagery examples of colonies for which earliest analyzed imagery predates known studies or reports. Earliest imagery shown on the left side and modern imagery shown on the right.

For both visible-light and SWIR/NIR conditions, we observed a recurring pattern with the earliest imaging dates (September to November) containing a smaller and relatively stable guano patch that contrasts against the environment (S13). The patch expanded substantially up to several times its initial size from November onwards to the end of the season (S12). Visible-light measurements increase in size over the entire season, while SWIR/NIR measurements often decrease or disappear completely in the last images with frozen colony site surface (end-of-season, before sea ice melts), and many last imaging dates with frozen surface (in January/February) contain no SWIR/NIR response while still containing significant guano patches in visible light (S21).

We found that guano area was related to colony size when aggregated over the breeding season (Figure 3C). Using annual average guano area estimates derived from visible-band satellite imagery, we observed a significant positive association with chick counts from ground surveys (Spearman’s rank correlation ρ = 0.59, p = 0.017) - larger colonies tend to produce more extensive guano stains during the breeding season.

#### Modelled Seasonal and Long-Term Patterns in Guano Area

The GAM model predicting intra-annual and inter-annual variation in guano area explained 63% of the deviance (adjusted R^2^ = 0.62) with highly significant effects for both day of year (edf = 5.46, F = 51.1, p < 0.001) and decimal date (edf = 4.88, F = 15.0, p < 0.001). No significant difference was found between VIS and SWIR sources (p = 0.61), supporting the robustness of the trend across sensor types. A separate model replacing the smooth function of decimal date with a linear term showed no evidence of a monotonic temporal trend (p = 0.61), suggesting that interannual variation in guano area is better captured by a flexible non-linear function than by a consistent increase or decrease through time. The model predicted a seasonal low of ∼22,000 m^2^ near DOY 90 (late austral summer), and a seasonal high of >1.7 million m^2^ near DOY 365 and 1 (early austral summer and late spring) (Figure 3). The long-term trend showed a non-linear pattern with fluctuations in total guano area that were not strictly monotonic. Despite occasional observational uncertainty from image artifacts, visual inspection and weighted modeling suggest that temporal variation reflects ecological signals rather than methodological noise. Predicted trends were consistent across data sources and confirmed through inspection of fitted curves with 95% confidence intervals (Figure 3).

## Discussion

We integrated historical datasets for detecting emperor penguin colonies and tracking their temporal change, using guano area as a proxy for colony size. Our study highlights the value of integrating historical and modern remote sensing datasets with field based surveys to expand species knowledge (e.g., highlighted in M. LaRue et al., 2024), including the temporal range and density of colony detections across the Antarctic. For 18 colonies, we provide the earliest confirmation of emperor penguin presence at or around the contemporary colony site. Furthermore, we show that the Cape Washington colony seems to have changed little in its size (as indicated by the guano patch) since 1972, despite ongoing global changes and relatively large interannual swings in adult counts at the colony (Kooyman & Ponganis, 2017). Taken together, these results suggest an impressive resilience of the species to changing environmental conditions over long time periods.

Historical Landsat sensors (MSS, TM, ETM+) detected guano prior to 2003, and as far back as 1972. Landsat MSS, TM, ETM+, OLI and Sentinel-2 datasets are suitable to track guano area change over time, as well as for mapping differences in guano spread over the breeding season. Our study found applicability of Keyhole imagery only for colonies that have high resolution data available and are relatively large in size, such as Cape Washington. Our guano area estimation showed moderate correlation with colony size, and emphasizes the need for similar studies covering multiple colonies.

Historical datasets enabled us to extend the reported colony presence back in time for several colonies. Keyhole imagery, especially high-resolution sensors such as KH-9 HEXAGON, can be used for detecting guano patches, as is exemplified with Cape Washington. Furthermore, the oldest Landsat MSS sensor datasets were consistent in their revisit time and coverage, and the georeferenced and digitally-stored data of four spectral bands made guano detection and discrimination from the environment easier than from Keyhole imagery, which has no spatial reference, suffers from digitization errors, and contains only a single panchromatic band. Newer Landsat sensors, such as ETM+, are further enhancing this detection power by including short-wave infrared bands (USGS, 2024).

Our analysis highlighted limitations of the Keyhole data - the irregular, scarce revisit time over the Antarctic and a single panchromatic band prevent detailed discrimination of guano patches. Older Keyhole sensors (KH-1/4 Corona and KH-5 Argon) have prohibitively coarse resolution (140 m) (USGS, 2008), and the images have a high degree of geometric and radiometric errors (Dashora et al., 2007) due to weather conditions and imperfect data recording, storage, and digitization (Fowler, 2004). Taken together, these limitations impede long-term monitoring or timeline reconstruction from the Keyhole dataset alone, but the data can instead serve as momentary observations of a colony. However, we caution that the Keyhole data available for download or ordering on USGS represents only a fraction of the entire produced dataset (Munteanu et al., 2024). Such data as well as historical aerial surveys, represent valuable resources for research if made available (Munteanu et al., 2024). Landsat data was more reliable, but images are still affected by unfavorable climatic conditions and technical errors such as cloud cover, SWIR noise, and SLC-off issues (applicable to Landsat 7), which limit the amount of usable imagery.

Although for many colonies, the Keyhole and Landsat datasets offer only complementary presence information, we identified the earliest detected guano patches which predate colony discovery for several colonies (Table 1). This is a novel addition to the long-term historical emperor penguin colony monitoring effort, and could be expanded to cover other areas across the Antarctic to search for abandoned colonies previously known to host penguins or to attempt to detect colonies in areas not yet known to researchers.

The guano area change analyses revealed changes within seasons and multi-annual trends between 1972 and 2024. Guano patch size increases modestly at the beginning of the breeding season, retaining a distinctly dense, dark and sharp-edged patch shape, but as the season progresses, the guano area expands and becomes more diffused in the environment (Lynch et al., 2012). These changes mirror emperor penguin breeding cycles, because the penguins forage more frequently and for longer as the season progresses, and then commence their annual departure from the colony for moulting (G. Kooyman et al., 2000; Wienecke, 2015). Our work supports evidence that guano diffuses and denudes as the season progresses (Lynch et al., 2012). Weathering- and mobility-induced guano changes result in the diffusion and spread of the remotely-sensed guano patch, and the formation of visible guano pathways away from the colony site. Moreover, an early decrease of SWIR/NIR guano response is visible, sometimes resulting in no guano response in SWIR/NIR for the last days or weeks of the breeding season, even though guano remains observable under visible light. The decrease in SWIR/NIR guano response was observed for images leading up to the thawing of the sea ice at the colony site in late December or January. This drop could be the result of organic decomposition affecting the reflectivity in the NIR and SWIR, as was previously noted in similar studies using cattle manure (Ben-Dor, 1997), and could be further modulated by local conditions such as denudation, weather and noise in the imagery.

Our long term analyses of guano change revealed a significant positive correlation of guano area coverage and colony size for the Cape Washington colony. While our data had some limitations, most of these were mitigated by our modelling approach. For instance, in ∼63% of all images, the guano patch was shadowed by topography, which may also induce measurement errors, and in SLC-off ETM+ imagery (post-Line Corrector failure (U.S. Geological Survey, 2003)), data gaps (present in 60% of all used ETM+ images) limit area estimate accuracy. To mitigate such errors, our models accounted for weighted minimum and maximum area estimates. Overall, our area estimates can be considered conservative.

Historical remote sensing imagery such as legacy Landsat (MSS, TM, ETM+) or Keyhole can provide novel species presence information, suggesting that publicly available and well-known datasets are valuable for ecological and conservation research, yet underutilized. When analyzed at broad spatial scales beyond just the confirmed colony locations, the data has the potential of revealing the existence of undiscovered colonies, similar to high-resolution contemporary remote-sensing datasets. Such methods could detect the historical existence of unknown colonies or expand our knowledge of existing colonies. Great potential also lies in conducting similar analyses for all colonies across the Antarctic to refine observed patterns of this species. Novel data driven detection methods including deep learning (Barthelme et al., 2024) can contribute to a better understanding of emperor penguin population changes, breeding, and responses and adaptation to climate change. Confirmation of colony locations prior to known dates, corroborated with the overall surprising size stability of the Cape Washington Colony suggest a high degree of species resilience to changing environmental conditions over time.

Remote sensing imagery is a challenging yet rewarding addition to conservation science in the Southern Ocean (Larue & Knight, 2014) and extending the time series of species presence and behavior with historic data helps contextualize species responses at local to regional scales.

Beyond advancing our fundamental understanding of a large marine bird, historic context can also facilitate conservation and management decisions at many scales (Munteanu et al., 2022, 2024; Rizayeva et al., 2023). This study showed that underutilized imagery sources such as Keyhole have a limited use in penguin studies, but coupled with other historical datasets such as Landsat, they can provide insights into colony dynamics over time, potentially predating the oldest existing reports for several colonies (Ancel et al., 2014; Fretwell et al., 2012; Fretwell & Trathan, 2009, 2021; Hempel & Stonehouse, 1987; M. A. LaRue et al., 2015; Lea & Soper, 2005; Plötz et al., 1985; Wienecke, 2012). Remote sensing can be a valuable addition to ground surveys, especially at locations where field visits are limited or remoteness prevents them altogether. Remotely-sensed guano presence can also be used to estimate colony size if certain limitations, such as imagery scarcity or guano patch diffusion, can be mitigated. The improved understanding of emperor penguin colony dynamics supplied by remote sensing data can support the current efforts in the conservation of this noble Antarctic species.

## Supporting information

Bielinis et al. supplememtary info

## Acknowledgements

MB acknowledges support through the Erasmus+ Traineeship Grant programme. CM acknowledges support by the German Science Foundation (DFG), Research Training Group ConFoBi (GRK 2123/1 TPX). BMK acknowledges support through an Early Career Fellowship at the Freiburg Institute for Advanced Studies.

## Data Accessibility

The data gathered during this study is accessible on GitHub: https://github.com/MartynasBielinis/penguins-data/

## Authors’ contributions

MB and CM conceptualized the study; MB collected and analyzed the data; MB, MLR and CM wrote the first draft, BMK performed the statistical analyses. All authors interpreted the results, contributed to writing and revision.

