## Supplementary material for "Historical remote sensing highlights long-term persistence of Emperor Penguin (*Aptenodytes forsteri*) colonies": Bielinis et al. supplememtary info

**S1. Study colonies**

| **Colony Name** | **Colony ID** | **Latitude** | **Longitude** | **Also known as** |
| --- | --- | --- | --- | --- |
| Amanda Bay | AMAN | -69.271 | 76.835 |  |
| Amundsen Bay | AMUN | -66.783 | 50.544 |  |
| Astrid | ASTR | -69.948 | 8.318 | Astrid Ice Tongue |
| Atka | ATKA | -70.614 | -8.132 | Atka Bay |
| Auster | AUST | -67.397 | 63.974 |  |
| Barrier Bay | BARR | -67.225 | 81.931 |  |
| Bear Peninsula | BEAR | -74.392 | -110.192 |  |
| Beaufort Island | BEAU | -76.929 | 166.881 | Beaufort Island North |
| Bowman Islands | BOWM | -65.161 | 103.067 | Bowman Island |
| Brownson | BROW | -74.14 | -103.48 | Brownson Islands |
| Bryan Coast | BRYA | -73.249 | -85.348 |  |
| Burton Ice Shelf | BURT | -66.783 | 89.45 | Posadowsky Bay |
| Cape Colbeck | COLB | -77.14 | -157.7 | Edward VII Peninsula |
| Cape Crozier | CROZ | -77.46 | 169.32 |  |
| Cape Darlington | CPDT | -71.887 | -60.134 |  |
| Cape Darnley | DARN | -67.887 | 69.696 | Flutter |
| Cape Gates | CPGT | -73.661 | -122.697 |  |
| Cape Poinsett | POIN | -65.782 | 113.235 |  |
| Cape Roget | ROGE | -71.988 | 170.597 |  |
| Cape Washington | WASH | -74.637 | 165.382 |  |
| Casey Bay | CASE | -67.312 | 46.957 | Casey Bay West |
| Coulman Island | COUL | -73.348 | 169.624 |  |
| Cruzen Island | CRUZ | -74.734 | -140.327 |  |
| Davis Bay | DAVI | -69.348 | 158.492 | Davies Bay |
| Dawson | DAWS | -76.014 | -26.648 | Dawson-Lambton |
| Dibble Glacier | DIBB | -66 | 134.8 |  |
| Dion Islands | DION | -67.865 | -68.71 | Emperor Island |
| Dolleman | DOLL | -70.611 | -60.421 | Dolleman Island |
| Drescher | DRES | -72.826 | -19.326 | Drescher Inlet |
| Fold Island | FOLD | -67.324 | 59.316 |  |
| Franklin Island | FRAN | -76.187 | 168.44 |  |
| Gould | GOUL | -77.71 | -47.656 | Gould Bay |
| Gunnerus | GUNN | -68.763 | 34.382 | Gunnerus Peninsula |
| Halley | HALL | -75.54 | -27.43 | Halley Bay |
| Haswell Island | HASW | -66.531 | 93.008 |  |
| Karelin Bay | KARE | -66.412 | 85.384 |  |
| Kloa Point | KLOA | -66.641 | 57.278 |  |
| Larsen Ice Shelf | LARS | -66.1 | -60.674 | Jason Peninsula |
| Lazarev | LAZA | -69.75 | 15.549 | Lazarev Ice Shelf |
| Ledda Bay | LEDD | -74.42 | -130.96 |  |
| Luitpold | LUTI | -77.271 | -33.552 | Luitpold Coast |
| Mertz Breakoff | MBRE | -66.892 | 146.62 |  |
| Mertz Glacier | MGLA | -67.24 | 145.535 |  |
| Ninnis Bank | NINN | -66.723 | 149.677 |  |
| Noville Peninsula | NOVI | -71.769 | -98.447 |  |
| Peterson Bank | PETE | -65.918 | 110.236 |  |
| Pfrogner Point | PFPT | -72.569 | -89.906 |  |
| Point Geologie | GEOL | -66.674 | 140.005 |  |
| Porpoise Bay | PORP | -66.32 | 129.75 |  |
| Ragnhild | RAGN | -69.908 | 27.155 | Ragnhild Coast |
| Riiser | RIIS | -72.124 | -15.106 | Riiser-Larsen |
| Rothschild | ROTS | -69.521 | -72.229 | Rothschild Island |
| Rupert Coast | RUPE | -75.382 | -143.308 |  |
| Sabrina Coast | SABR | -66.177 | 121.058 |  |
| Sanae | SANA | -69.999 | -1.413 |  |
| Shackleton Ice Shelf | SHAC | -65.089 | 96.02 |  |
| Smith | SMIT | -74.369 | -60.827 | Smith Peninsula |
| Smyley | SMYL | -72.302 | -78.82 | Smyley Island |
| Snow Hill | SNOW | -64.524 | -57.444 | Snow Hill Island |
| Stancomb | STAN | -74.12 | -23.087 | Stancomb Wills Glacier |
| Taylor Glacier | TAYL | -67.454 | 60.878 |  |
| Thurston Glacier | THUR | -73.498 | -125.62 | Mount Siple |
| Umebosi | UMBE | -68.046 | 43.017 | Umbeashi |
| Verdi Inlet | VDIT | -71.556 | -74.76 |  |
| West Ice Shelf | WEST | -66.55 | 81.818 |  |
| Yule Bay | YULE | -70.716 | 166.478 |  |

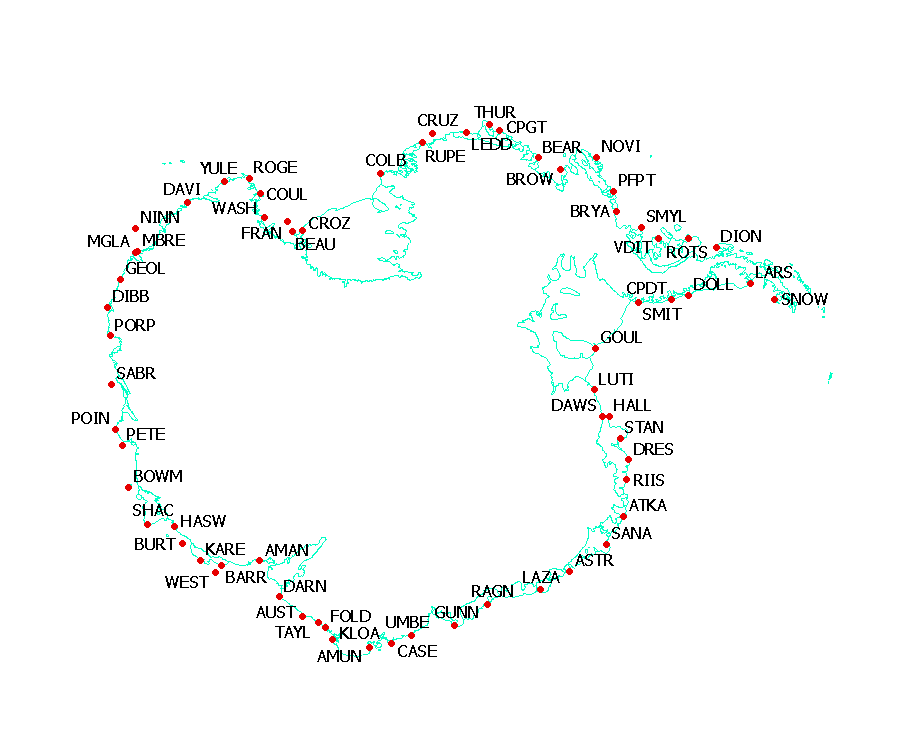

**S2**. Map of study colonies. Antarctic land and ice shelf boundaries pictured in cyan (Gerrish et al., 2020)

**S3. Keyhole sensor specifications as described by USGS**

| **Sensor Name** | **Mission Designator** | **Best Ground Resolution** | **Declass Dataset** | **Acquisition Period** |
| --- | --- | --- | --- | --- |
| CORONA | KH-1; KH-2; KH-3; KH-4 | 1.8 - 12 m | Declass-1 | 1960 - 1972 |
| ARGON | KH-5 | 140 m | Declass-1 | 1962 - 1964 |
| LANYARD | KH-6 | 1.8 m | Declass-1 | 1963 |
| GAMBIT | KH-7 | 0.6 - 1.2 m | Declass-2 | 1963 - 1967 |
| HEXAGON | KH-9 | 6 - 12 m | Declass-2 | 1973 - 1980 |
| HEXAGON | KH-9 | 6 - 12 m | Declass-3 | 1971 - 1984 |

**S4. Landsat mission specifications, as described by USGS. Note that Thermal IR and Panchromatic bands in TM/ETM+ are not utilized, and are captured at coarser resolution than specified here.**

| **Landsat Mission** | **Sensor** | **Best Ground Resolution** | **Bands** | **Acquisition Period** |
| --- | --- | --- | --- | --- |
| 1, 2, 3, 4, 5 | Multispectral Scanner (MSS) | 80 m (processed to 60 m) | 4: Green, Red, 2x NIR | 1972 - 1992 |
| 4, 5 | Thematic Mapper (TM) | 30 m | 7: Blue, Green, Red, NIR, 2x SWIR, Thermal IR | 1982 - 2012 |
| 7 | Enhanced Thematic Mapper (ETM+) | 30 m | 7: Blue, Green, Red, NIR, 2x SWIR, Thermal IR, Panchromatic | 1999 - 2024 (SLC-off since 2003) |

**S5. Sensor specifications of additional imagery sources for guano area measurement**

| **Mission** | **Sensor** | **Best Ground Resolution** | **Bands** | **Acquisition Period** |
| --- | --- | --- | --- | --- |
| Landsat-8/9 | Operational Land Imager (OLI) | 30 m | 9: Coastal Aerosol, Blue, Green, Red, NIR, 2x SWIR, Panchromatic, Cirrus | 2013 - now |
| Sentinel-2 | Multispectral Instrument (MSI) | 10 m | 13: Aerosol, Blue, Green, Red, 4x Red Edge, NIR, Water Vapor, 3x SWIR | 2015 - now |

**S6. Acquired imagery for historical colony detection**

| **Sensor/Dataset** | **Acquisition Period** | **Filtering Parameters** | **Images Acquired** |
| --- | --- | --- | --- |
| Keyhole Imagery | Dec 1974 – Sep 1980 | Declass-2 and Declass-3 imagery | 44 |
| Landsat 1-5 (MSS) | Nov 1972 – Jan 1974 | All imagery | 146 |
| Landsat 4-5 (TM) | Dec 1988 – Feb 1992 | All imagery | 128 |
| Landsat 7 (ETM+) | Jan 2000 – May 2003 | All SLC-on data | 202 |

**S7. Acquired imagery for guano area measurement**

| **Sensor/Dataset** | **Acquisition Period** | **Filtering Parameters** | **Images Acquired** |
| --- | --- | --- | --- |
| Keyhole Imagery | Dec 1974 – Sep 1980 | High-resolution imagery only | 5 |
| Landsat 1-5 (MSS) | Nov 1972 – Jan 1974 | All imagery | 10 |
| Landsat 4-5 (TM) | Dec 1988 – Feb 1992 | All imagery | 15 |
| Landsat 7 (ETM+) | Jan 2000 – Feb 2013 | All SLC-on data (smaller set of imagery), then SLC-off data every ~6 days | 104 |
| Landsat 8-9 (OLI) | Oct 2013 – Jan 2016 | Every ~6 days. | 37 |
| Sentinel-2 | Oct 2016 – Feb 2024 | Every ~6 days | 86 |

**S8. Highest colony coordinate offsets between the original list and Fretwell study list** **(Fretwell & Trathan, 2021)** **(>100m).**

| **Colony Name** | **Offset** |
| --- | --- |
| West Ice Shelf | 76.88 km (location corresponding with Barrier Bay) |
| Barrier Bay | 74.87 km (location corresponding with West Ice Shelf) |
| Brownson | 23.99 km |
| Ledda Bay | 18.59 km |
| Cape Colbeck | 4.97 km |
| Pfogner Point | 0.12 km |
| Verdi Inlet | 0.11 km |

**S9. Highest colony coordinate offsets between the original list and MAPPPD database (>100 m)**

| **Colony Name (original list)** | **Colony Name (MAPPPD)** | **Offset** |
| --- | --- | --- |
| Mertz Breakoff | Mertz Glacier | 62.88 km |
| Burton Ice Shelf | Posadowsky Bay | 58.03 km |
| Casey Bay | Casey Bay West | 15.50 km |
| Cape Colbeck | Cape Colbeck / Edward VII Peninsula | 5.52 km |
| Bear Peninsula | Bear Peninsula | 4.95 km |
| Beaufort Island | Beaufort Island North | 4.12 km |
| Halley | Halley | 1.69 km |
| West Ice Shelf | West Ice Shelf | 1.44 km |
| Cruzen Island | Cruzen Island | 1.42 km |
| Cape Crozier | Cape Crozier | 0.63 km |
| Barrier Bay | Barrier Bay | 0.57 km |
| Dion Islands | Emperor Island | 0.28 km |

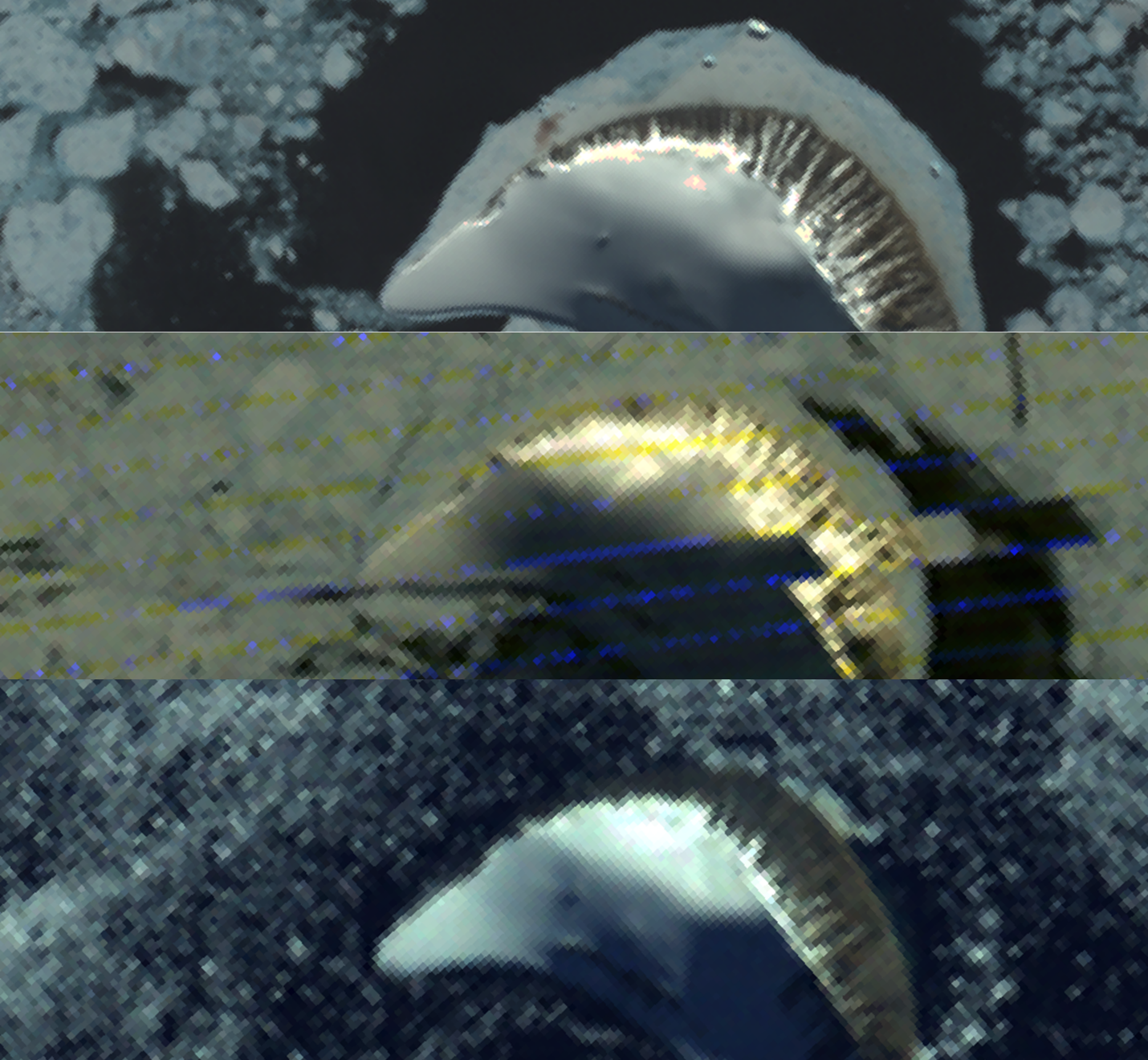

**S10**. Beaufort Island colony guano presence example. Top: Unambiguous (confirmed) guano presence (Landsat 7 ETM+, November 13^th^, 2000); Center: Uncertain/ambiguous presence (Landsat 2 MSS, October 25^th^, 1975); Bottom: No guano presence due to water (Landsat 1 MSS, January 5^th^, 1974). Note that lack of unambiguous guano presence might not mean colony absence.

**S11. Band combinations used in guano area measurements**

| **Sensor** | **Visible-light band combination** | **SWIR/NIR band combination** |
| --- | --- | --- |
| Sentinel-2 | B4-B3-B2 | B12-B11-B8a |
| Landsat OLI | B4-B3-B2 | B7-B6-B5 |
| Landsat ETM+ | B3-B2-B1 | B7-B5-B4 |
| Landsat TM | B3-B2-B1 | B7-B5-B4 |
| Landsat MSS | B6-B5-B4 | Not applicable |
| Keyhole | Single panchromatic band | Not applicable |

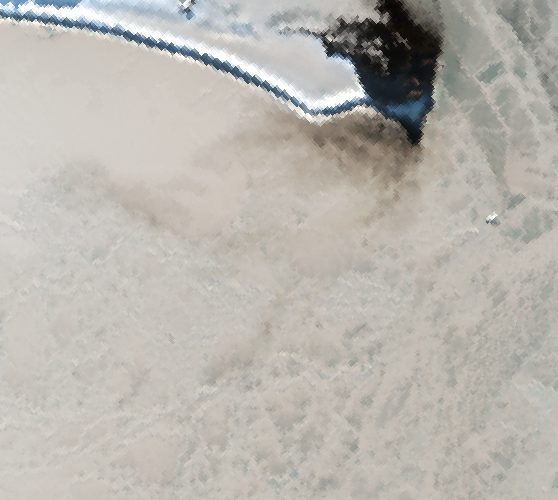

**S12**. Diffused guano patch with visible guano tracks (Sentinel-2, November 27^th^, 2019)

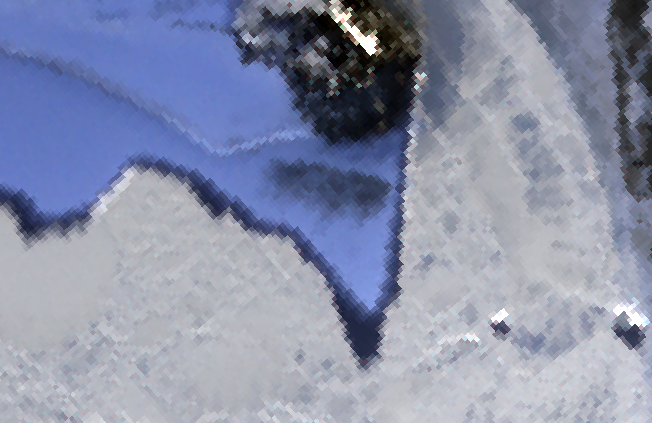

**S13**. Guano visible as a dense cluster (Sentinel-2, September 29^th^, 2019)

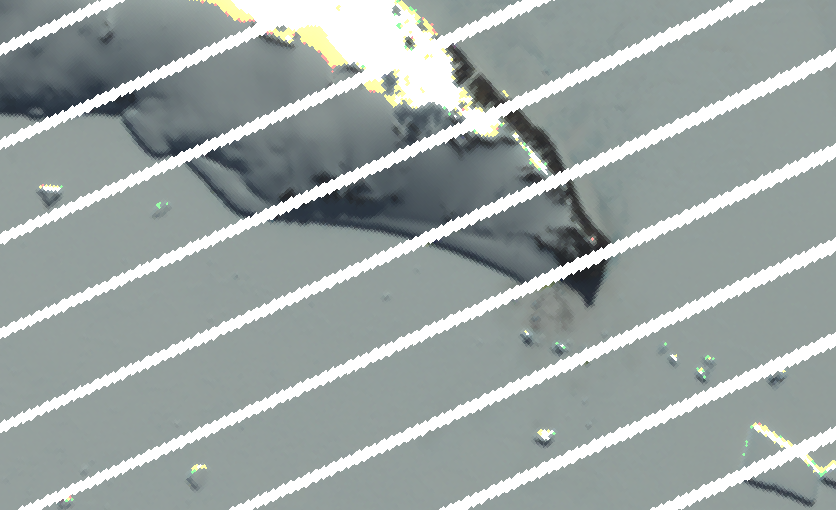

**S14**. Landsat 7 ETM+ data after the Scan Line Corrector (SLC) fault (SLC-off) in Cape Washington, December 15^th^, 2006.

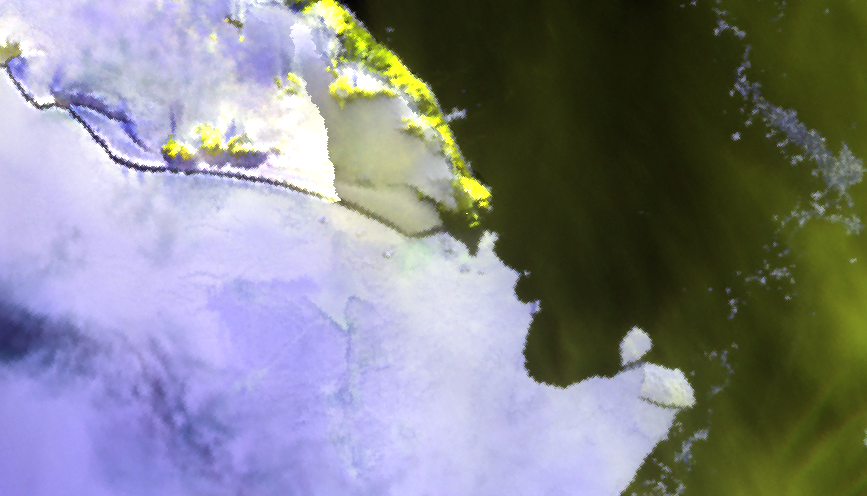

**S15**. Clouds as seen in short-wave infrared (yellow) Sentinel-2 data, obscuring the Cape Washington colony guano signature on December 4^th^, 2016.

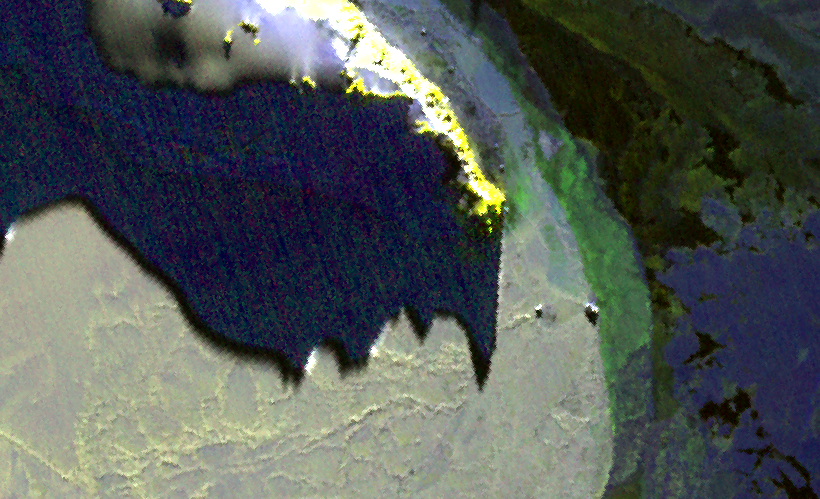

**S16**. High noise level and shadow obscuring possible colony guano signature in short-wave infrared (yellow) Sentinel-2 data at Cape Washington on September 19^th^, 2019.

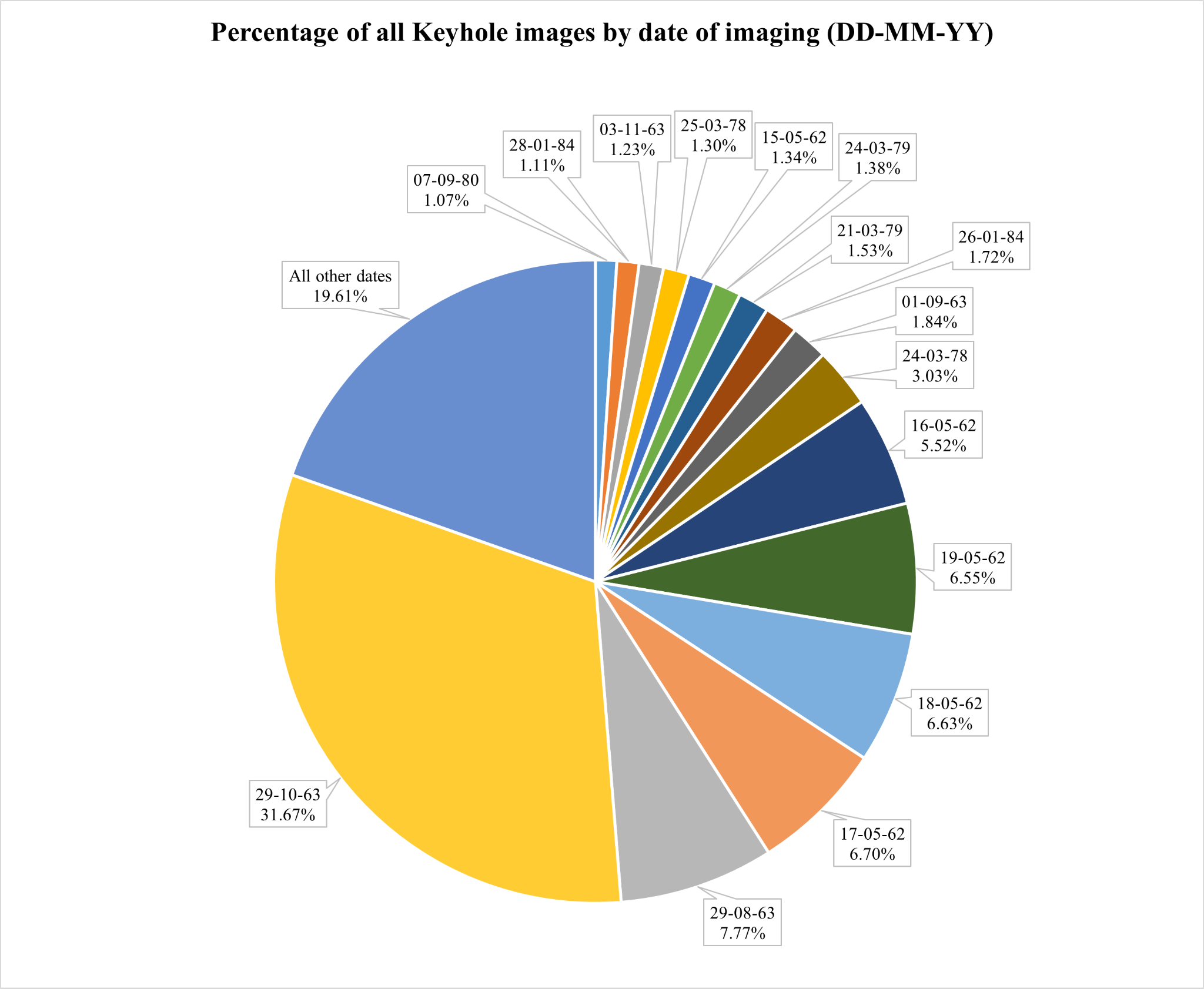

**S17**. Percentage of all Keyhole images by their date of imaging, including repeat uses of the same acquisition for multiple colonies.

**S18. Colonies with imagery from each Keyhole dataset**

| **Dataset** | **Colonies with Imagery** |
| --- | --- |
| Declass-1 | 66 of 66 |
| Declass-2 | 27 of 66 |
| Declass-3 | 30 of 66 |

**S19. Keyhole image availability**

| **Dataset** | **Average Number of Images per Colony** | **% of Download-ready Images** |
| --- | --- | --- |
| Declass-1 | ~29 | 70.8 |
| Declass-2 | ~6 | 37.67 |
| Declass-3 | ~17 | 22.09 |

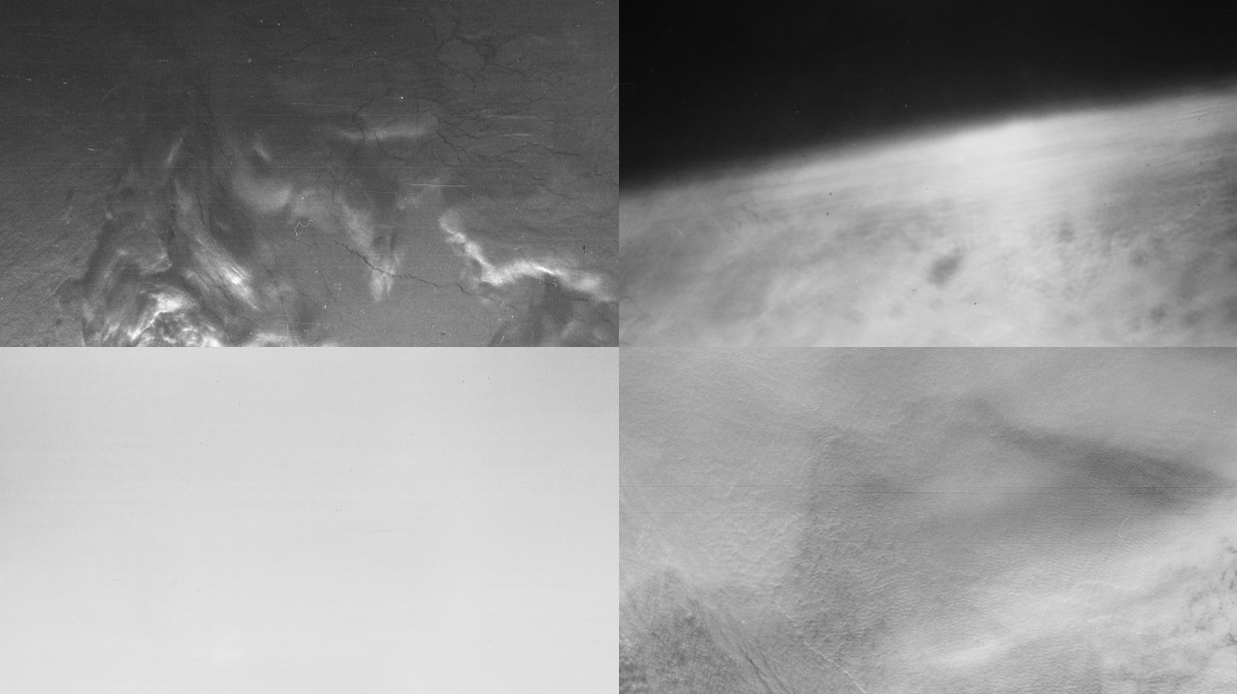

**S20**. Examples of unsuitable Keyhole imagery. From top right clockwise: Cloud cover and low brightness; overbrightened without surface visibility; complete cloud cover; over-brightened with possible cloud cover

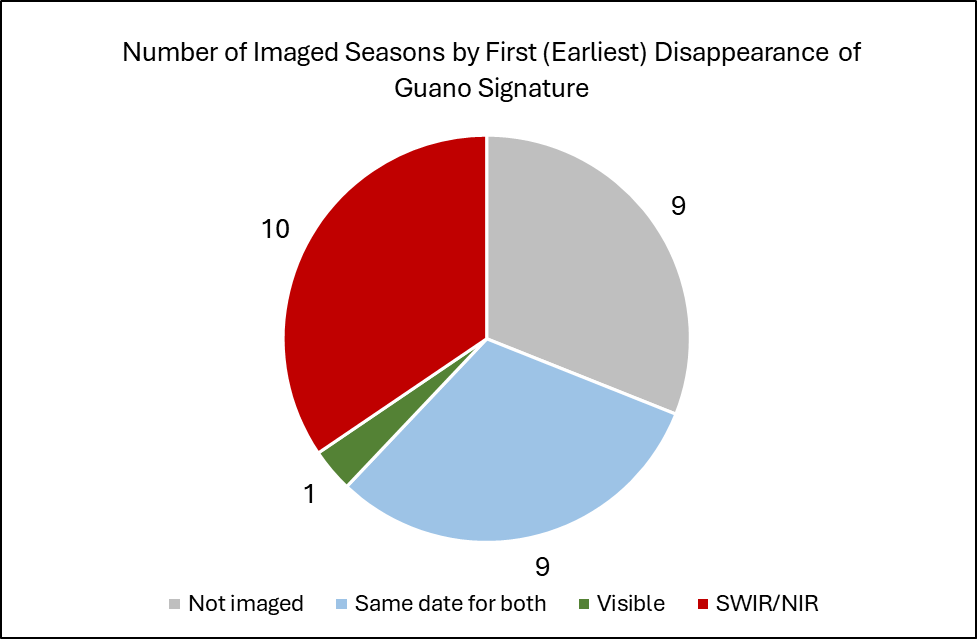

**S21**. Number of imaged seasons by first (earliest) disappearance of guano signature. Not imaged – guano disappearance is not imaged; Same date for both – guano patch disappears in both band combinations on the same date; Visible – guano disappears in visible-light before SWIR/NIR; SWIR/NIR – guano disappears in SWIR/NIR combination before visible-light.

**Supplementary Material References:**

Fretwell, P. T., & Trathan, P. N. (2021). Discovery of new colonies by Sentinel2 reveals good and bad news for emperor penguins. *Remote Sensing in Ecology and Conservation*, *7*(2), 139–153. https://doi.org/10.1002/rse2.176

[Gerrish, L., Fretwell, P., & Cooper, P. (2020). *Medium resolution vector polygons of the Antarctic coastline* (Version 7.3, p. 2 files, 7 MB) [Http://www.opengis.net/wms,application/geopackage+sqlite3,application/xml]. UK Polar Data Centre, Natural Environment Research Council, UK Research & Innovation. https://doi.org/10.5285/ED0A7B70-5ADC-4C1E-8D8A-0BB5EE659D18](https://www.zotero.org/google-docs/?n47xe9)
